## Supplemental Information for "Top-down modulation of sensory processing and mismatch in the mouse posterior parietal cortex"

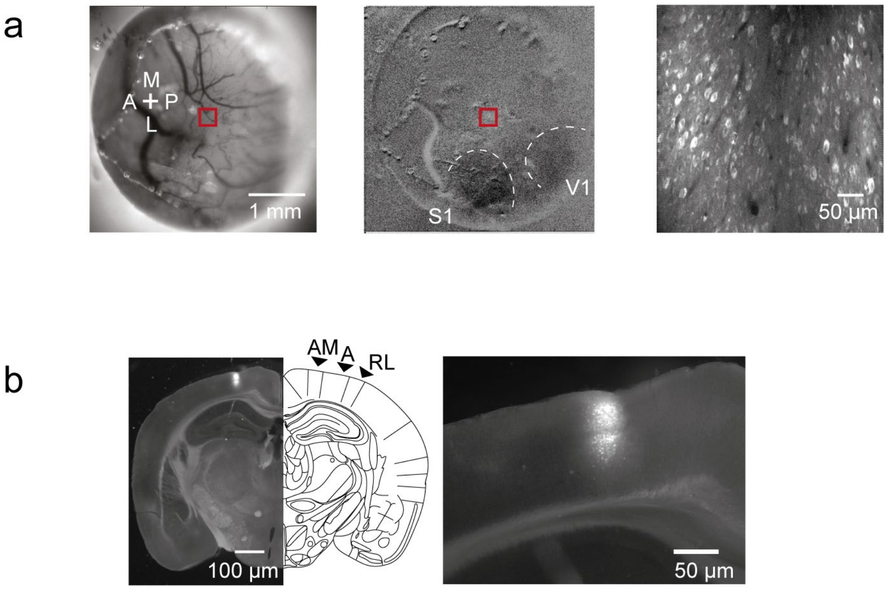


**Supplementary Fig. 1. Locating area A of the PPC through intrinsic optical signal imaging (IOS).** **a** Left, view of a chronically implanted cranial window over the PPC, with the imaging site outlined in red. Center, location of the γ and δ -whisker barrel in S1 and of V1, as determined with IOS seen through a 4-mm cranial window. Right, view of the imaged field of view (red box) of layer 2/3 neurons expressing the GECI RCaMP1.07, as acquired with the 2-photon microscope. **b** Left, transverse section of a brain slice with RCaMP1.07 expression in layers 2/3 of the PPC. Right, a close-up of the GECI expression site.

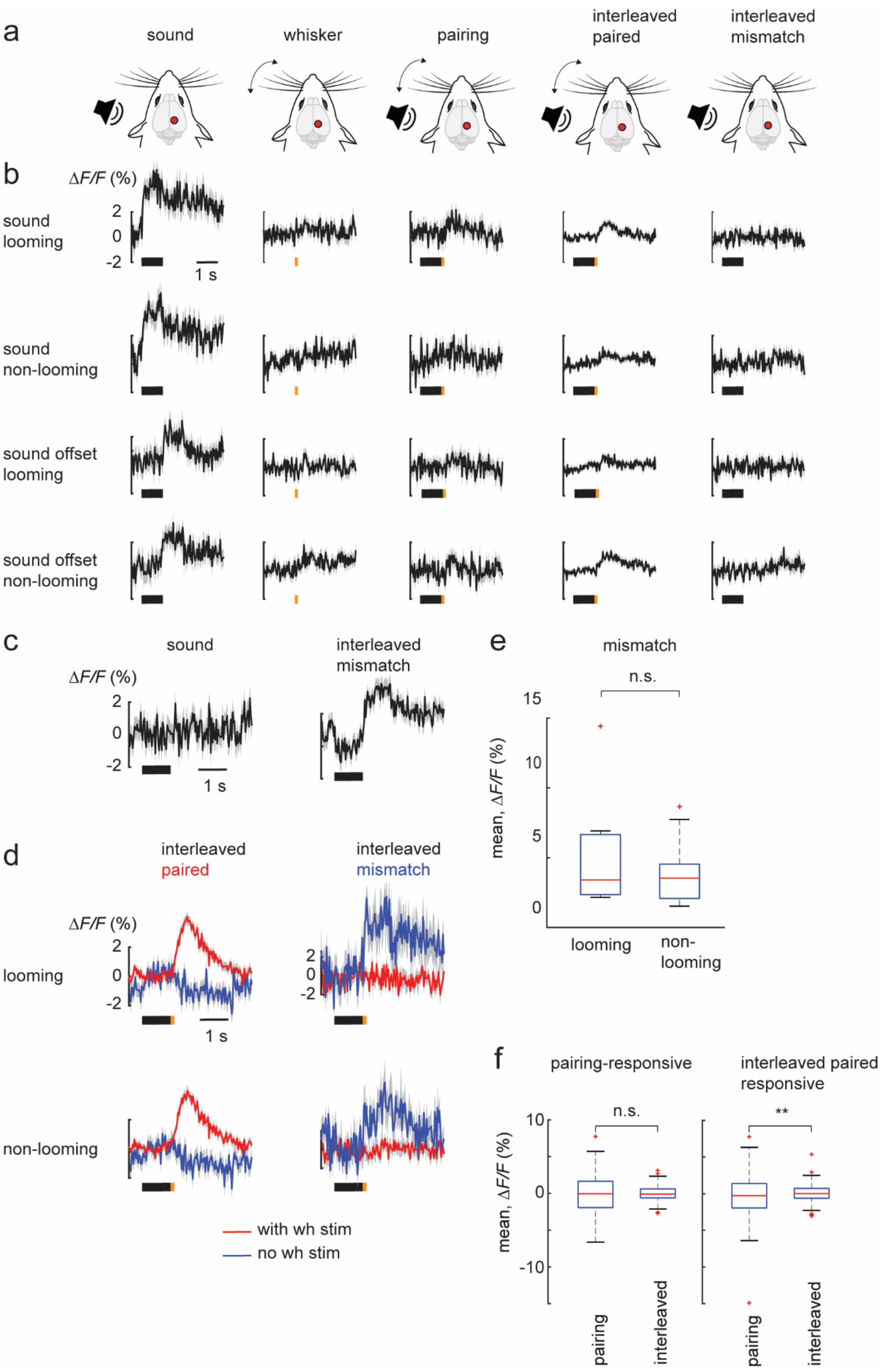


**Supplementary Fig. 2. Creating a sensory association at the PPC.**

**a** Schematic of the sequence of stimulus presentation (same as Fig. 1a). **b** Same as **Fig. 1d**, for looming sound (104 neurons), non-looming sound (101 neurons), looming sound offset-responsive (n = 100) and non-looming sound offset-responsive (117 neurons) neurons. **c** Average population responses of mismatch-responsive neurons (from Fig. 1d) in their corresponding looming sound trials. Note the absence of a sound offset response in the sound trials for these neurons. **d** Population averages (± s.e.m.) of ∆F/F traces of interleaved paired and mismatch-responsive neurons when the whisker stimulus is associated with a looming and non-looming sound respectively. **e** Boxplots of average population responses of mismatch-responsive neurons, with a looming (9 neurons) and non-looming (11 neurons) sound. wh stim: whisker stimulus. n.s. refers to non-significant here and in all subsequent figures. **f** Boxplot from Fig. 1g with outliers. Data represented as mean ± s.e.m.

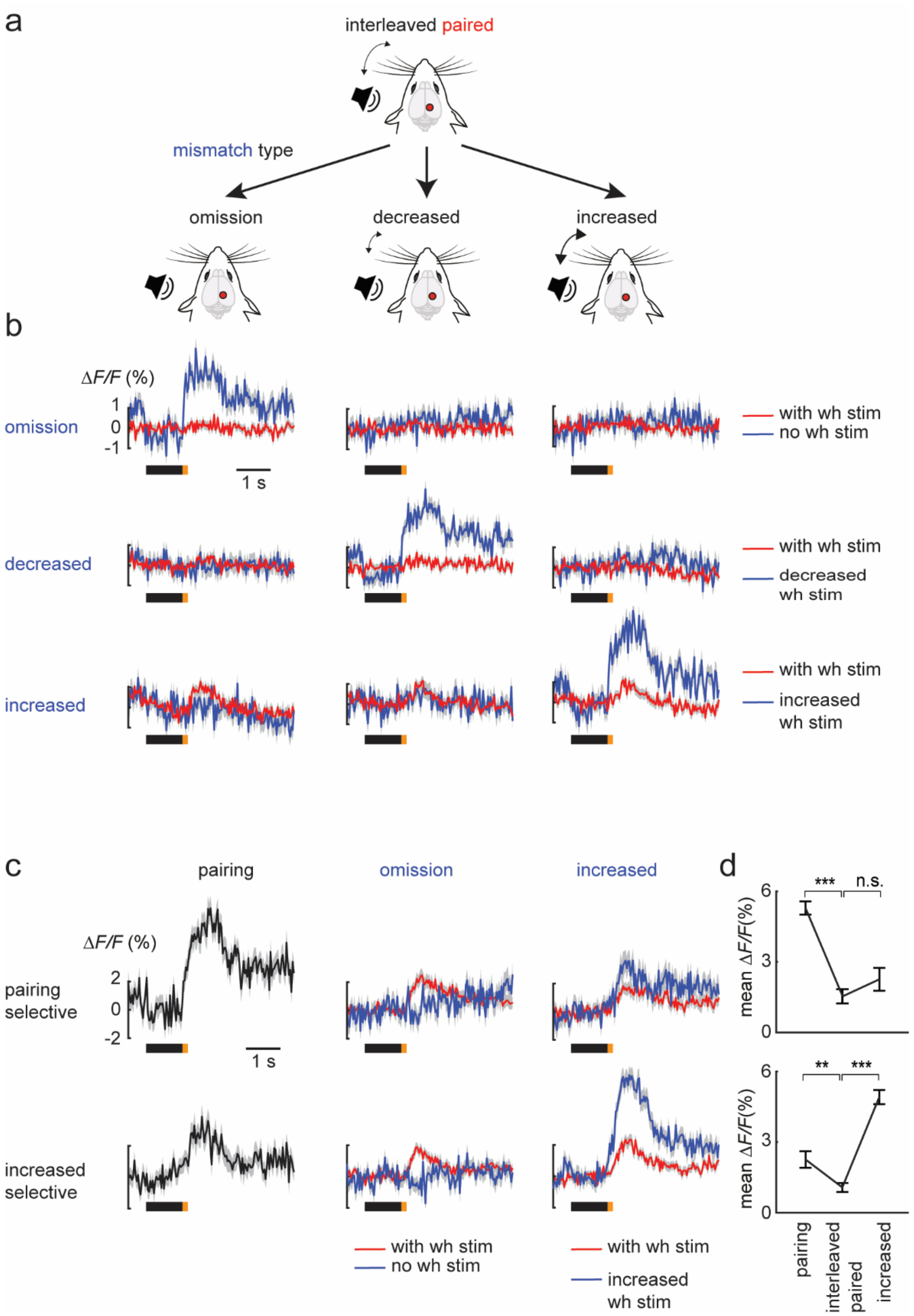


**Supplementary Fig. 3. PPC neurons can report different types of mismatch.**

**a**-**b** Same as **Fig. 3e-f**, with corresponding population average traces across the different mismatch trials for each mismatch selective population. **C** Population averages (± s.e.m.) of ΔF/F traces of pairing (61 neurons) and increased-responsive (mismatch) neurons (107 neurons), across the pairing, omission-mismatch and increased-mismatch sessions. **d** Average population responses of the neurons in **c** across the sessions. wh stim: whisker stimulus. Data represented as mean ± s.e.m. Statistical significance is indicated by * for $p < 0.05$, ** for $p < 0.01$, and *** for $p < 0.001$, with two-sided Wilcoxon signed-rank paired test.

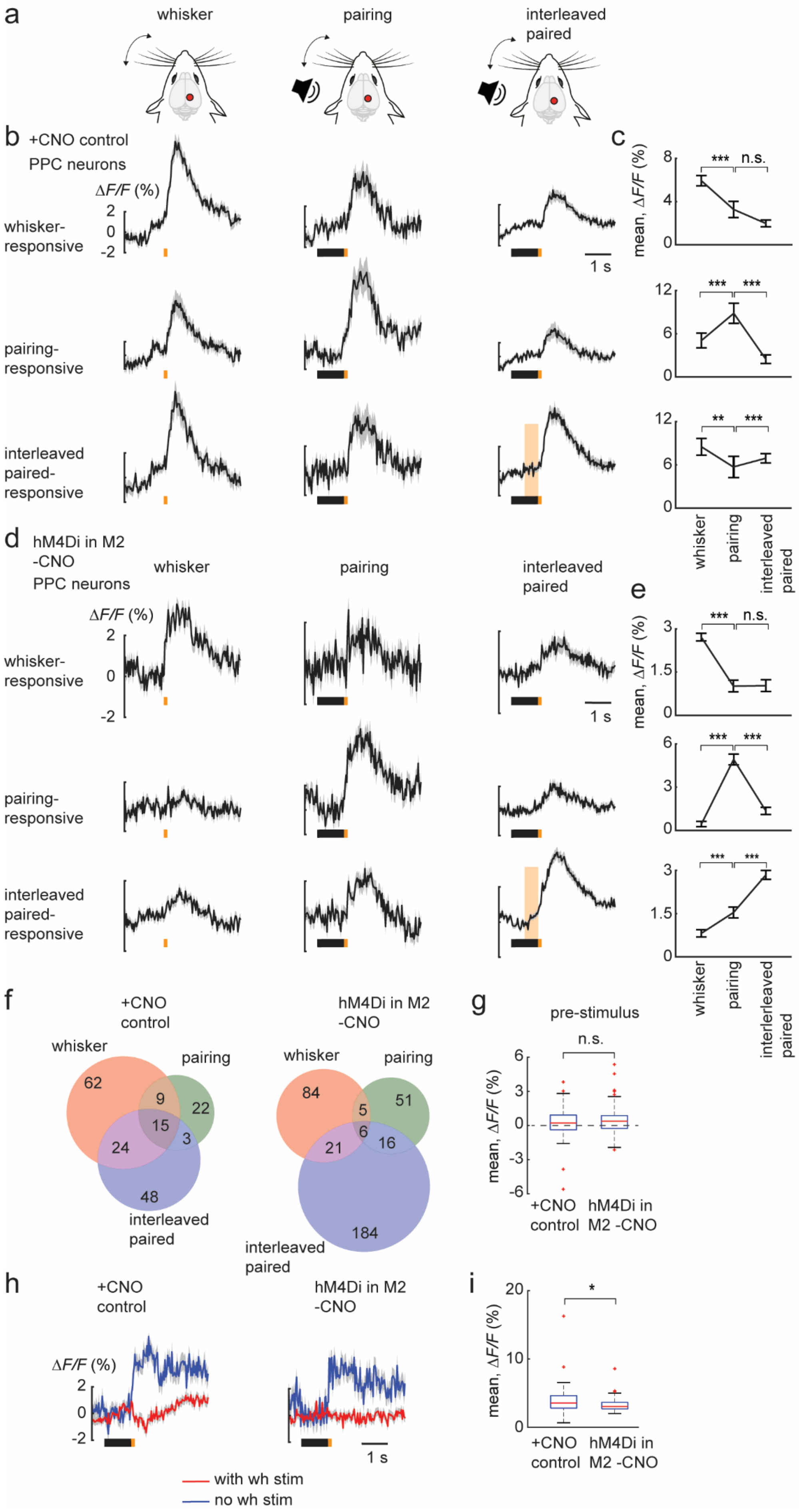

a
whisker
pairing
interleaved
paired
b +CNO control
PPC neurons
ΔF/F (%)
2
0
-2
whisker-
responsive
pairing-
responsive
interleaved
paired-
responsive
1 s
c
mean, ΔF/F (%)
8
4
0
***
n.s.
12
6
0
***
***
12
6
0
**
***
whisker
pairing
interleaved
paired
d hM4Di in M2
-CNO
PPC neurons
whisker
pairing
interleaved
paired
ΔF/F (%)
2
0
-2
whisker-
responsive
pairing-
responsive
interleaved
paired-
responsive
1 s
e
mean, ΔF/F (%)
3
1.5
0
***
n.s.
6
3
0
***
***
3
1.5
0
***
***
whisker
pairing
interleaved
paired
f
+CNO
control
whisker
pairing
62
9
22
15
24
3
48
interleaved
paired
hM4Di in M2
-CNO
whisker
pairing
84
5
51
6
21
16
184
interleaved
paired
g
pre-stimulus
n.s.
mean, ΔF/F (%)
6
3
0
-3
-6
+CNO
control
hM4Di in
M2 -CNO
h
+CNO
control
hM4Di in M2
-CNO
ΔF/F (%)
2
0
-2
1 s
with wh stim
no wh stim
i
20
10
0
*
mean, ΔF/F (%)
+CNO
control
hM4Di in
M2 -CNO

**Supplementary Fig. 4. Effects of CNO on sensory processing.**

**a** Schematic of the sequence of stimulus presentation (same as **Fig. 2a**). **b-e** Populaton averages (± s.e.m.) of Δ*F/F* traces of whisker-, pairing- and paired-responsive neurons in layer 2/3 of the PPC, along with their corresponding population average for the other sessions +CNO (4 mice, 6 FOVs) and without CNO (3 mice, 6 FOVs). Neuron numbers of the respective sessions are shown in the Venn diagrams in **f**. **f** Venn diagrams of session-responsive neurons in **b**,**d**, with their respective overlaps between sessions. **g** Box plots of pre-stimulus response of interleaved paired-responsive neurons +CNO (90 neurons) and without CNO (227 neurons). **h** Population averages (± s.e.m.) of ΔF/F traces of interleaved mismatch-responsive neurons + CNO and without CNO. **i** Box plots of average population responses of mismatch-responsive neurons +CNO (78 neurons) and without CNO (57 neurons). wh stim: whisker stimulus. Data represented as mean ± s.e.m. Statistical significance is indicated by *** for $p < 0.001$, with two-sided Wilcoxon signed-rank paired test (**c**,**e**) and two-sided Wilcoxon-Mann-Whitney test (**g**, **i**).

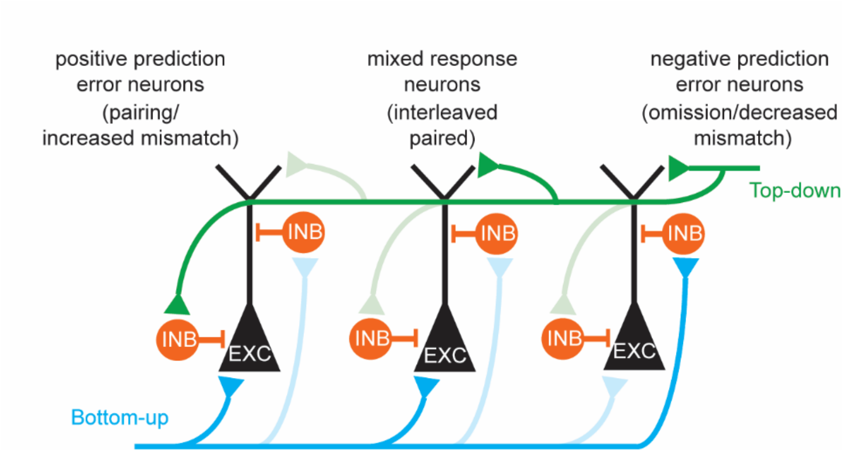


**Supplementary Fig. 5. Microcircuit for predictive processing in the PPC.**

We have adapted the proposed canonical microcircuit for predictive processing to include the respective session-responsive neurons that we have identified, in this simplified model restricted to layer 2/3 neurons. In this model, the positive prediction error neurons receive bottom-up input in the form of the whisker stimulus, which is balanced by top-down inhibition. The pairing-responsive neurons could contribute to part of this class of positive prediction error neurons. They are suppressed in transition from the pairing to interleaved session, as the prediction is formed. These pairing-responsive neurons can than report the increased-mismatch (Supplementary **Fig. 3c-d**), where the whisker stimulus is stronger than expected, compared to the interleaved paired trials (positive prediction error). The interleaved paired-responsive neurons increase their response to the whisker stimulus in transition from the pairing to the interleaved session, as they now receive more top-down feedback/prediction (matched with an increased pre-stimulus response in **Fig. 2h**). Lastly the negative prediction error neurons receive top-down feedback that is balanced by bottom-up inhibition. Both the omission and decreased intensity mismatch-responsive neurons could represent these negative prediction error neurons as they are recruited during the omission of the whisker stimulus and the presentation of a decreased whisker stimulus intensity respectively.
